# The Class III peroxidase OsPrx20 is a key regulator of stress response and growth in rice

**DOI:** 10.1101/2024.04.08.588571

**Authors:** Tao Shen, Qingwen Wang, Dan Chen, Huining Ju, Runjiao Yan, Fengjuan Xu, Donghuan Fu, Xiaona Bu, Huan Zhang, Jiexiong Hu, Zhengguang Zhang, Lan Ni, Mingyi Jiang

## Abstract

In plants, reactive oxygen species maintain strictly low intracellular levels that is prerequisite for their function as second messengers. However, abiotic and biotic stresses are also long-term processes, requiring rapid scavenging of excess intracellular ROS. Plant Class III peroxidases, as multifunctional enzymes, are core enzymes in regulating intracellular ROS homeostasis and are key to regulating the complicated signaling network. Here, we found a rice Class III peroxidase OsPrx20 maintains intracellular ROS homeostasis in different conditions. OsPrx20 is a target of Ca^2+^/calmodulin-dependent protein kinase (OsDMI3) and phosphorylates its Thr-244 site by OsDMI3. Overexpression of *OsPrx20* enhances osmotic stress tolerance but reduces blast resistance, while lack of *OsPrx20* is opposite, suggesting OsPrx20 positively regulates osmotic stress tolerance but negatively regulates blast resistance in rice. Meanwhile, overexpression of *OsPrx20* enlarges spike size and grain fullness, whereas lack of *OsPrx20* leads to dwarfism and smaller spike. In ABA signaling, OsPrx20 Thr-244 phosphorylation is specifically dependent on OsDMI3 to reduce the sensitivity of ABA to seed germination and root growth and to enhance osmotic stress tolerance without affecting spike and grain development. Our study reveals an essential regulatory mechanism that directly activates OsPrx20 in ABA signaling, highlighting the multi-functionality of OsPrx20 in different biological processes.

## 1. Introduction

Rice, a sessile organism, growth is mainly threatened by environmental stimuli (cold, heat, drought, salt, and heavy metals) and diseases (rice blast, bacterial blight, and sheath blight). It has evolved a wide range of signal response mechanisms to adapt to these challenges. Reactive oxygen species (ROS), one of these signal molecules, is indispensable for plant metabolism and has emerged as a central signal molecule **(Apel and Hirt, 2004)**. Their roles, which act as signaling molecules that regulate growth and respond to environmental stimuli in plants, are widely recognized **(Baxter et al., 2014; Mittler et al., 2011; Wang et al., 2023a)**. The apoplast is an interface for exchanging nutrients and signals between plant cells and the environment. In most cases, plant responses to environmental and endogenous stimuli involve the accumulation of ROS within it **(Kimura et al., 2017)**. Apoplastic H_2_O_2_ production primarily results from plasma membrane-localized NADPH oxidases (respiratory burst oxidase homologs, RBOHs) **(Suzuki et al., 2011)**. In Arabidopsis, AtRbohD and AtRbohF promote H_2_O_2_ production to enhance salt-induced antioxidant defense **(Ben Rejeb et al., 2015)**. AtRbohD stimulates ROS production to regulate plant immunity **(Lee et al., 2020; Pfeilmeier et al., 2021)**. AtRbohC, AtRbohH, and AtRbohJ are involved in the polar growth of root hair through auxin regulation of ROS homeostasis **(Mangano et al., 2017)**. In rice, OsRbohB promotes apoplastic H_2_O_2_ production in response to abiotic stress **(Shi et al., 2020; Wang et al., 2023)**. Mutation of different *OsRbohs* leads to decreased rice blast, bacterial blight, and sheath blight resistance, and all *osrboh* mutants display one or more compromised yield traits **(Zhu et al., 2024)**. Indeed, even under stress, the intracellular ROS levels are tightly controlled; an increase in ROS production rates does not necessarily translate into elevated ROS homeostasis concentration **(Waszczak et al., 2018)**. Intracellular lifetimes of different ROS vary widely, from nanoseconds for ^•^OH, microseconds for ^1^O_2_, and milliseconds to seconds for O_2_^•-^, and H_2_O_2_, but the elevated ROS concentrations in separate subcellular compartments are all short-lived **(Mattila et al., 2015)**. However, either abiotic or biotic stress is a long-term process that provokes ROS homeostasis imbalance, triggering ROS accumulation **(Cejudo et al., 2021)**. Excessive intracellular ROS leads to oxidative damage, such as DNA and RNA damage, protein denaturation, enzyme inactivation, and membrane lipid peroxidation, resulting in cellular senescence or even death **(Mittler et al., 2022).** Therefore, plant cells possess a well-developed interlinked array of antioxidants to combat the threat of ROS. The enzymatic antioxidants consist of superoxide dismutase (SOD), catalase (CAT), ascorbate peroxidase (APX), and other related enzymes that can restrict ROS levels **(Bhatt and Tripathi, 2011; Mittler, 2002)**. In addition, a large family of peroxidases (EC 1.11.1.X; Prxs) in plants, which catalyze the detoxification of H_2_O_2_ and other peroxidases in different organisms, have also been identified.

Based on protein structure, the peroxidase superfamily can be classified as heme peroxidases (Hemo Prxs) and non-heme peroxidases (Non-hemo Prxs) **(Kidwai et al., 2020)**. Based on their functions, the heme Prxs family can be further divided into three subclasses: Class I (ascorbate peroxidases), Class II (lignin peroxidases), and Class III (secreted peroxidases), which share a similar three-dimensional structure and a heme group that is composed of protoporphyrin IX and Fe^3+^ **(Liu et al., 2021b)**. The Class I Prxs exist in most organisms except animals, especially prokaryotes, suggesting a possible origin for the other Prxs. They include cytochrome *c* peroxidases (EC 1.11.1.5; CcPs), catalase (EC 1.11.1.6; CAT), and ascorbate peroxidases (EC 1.11.1.11; APX), which have the primary function of the cell is to scavenge excess H_2_O_2_. The Class II Prxs include manganese peroxidases (EC 1.11.1.13; MnPs) and lignin peroxidases (EC 1.11.1.14; LiPs), which play a key role in the degradation of lignin-containing detritus in the soil, as none of the other peroxidases are capable of degrading lignin **(Cosio and Dunand, 2009; Jha et al., 2022)**. The Class III Prxs (EC 1.11.1.7), plant-specific heme oxidoreductases, are secreted into the extracellular space or vacuole, which are typically localized in the cytoplasm and cell wall, are highly thermally stable and naturally glycoproteins with a wide range of substrates and marked substrate specificity for phenols **(Cosio and Dunand, 2009; Shigeto and Tsutsumi, 2016)**. These characteristics clearly distinguish them from Class I Prxs, which are not glycoproteins and are mainly localized in chloroplasts, mitochondria, and peroxisomes. In addition, APX is substrate-specific for ascorbate, whereas Class III Prxs are not. Class III Prxs are widely distributed in the plant kingdom, with Arabidopsis (*Arabidopsis thaliana*) and rice (*Oryza sativa* L.) genomes encoding 73 and 138 Class III Prxs, respectively **(Liu et al., 2021b; Passardi et al., 2004; Zheng et al., 2023)**. They are involved in various physiological processes, including ROS production and cleavage, resistance to biotic and abiotic stresses, lignin biosynthesis, seed germination, and cell elongation. The role of Class III Prxs in defense and growth has been extensively explored in recent years. In the response of plants to abiotic stresses, there is a higher peroxidase activity by overexpression of Class III Prxs. For example, overexpression of *AtPrx22*, *AtPrx39*, and *AtPrx69* enhances cold tolerance in Arabidopsis **(Kim et al., 2012)**, overexpression of *TaPrx-2A* positively regulates salt stress tolerance by scavenging ROS in wheat **(Su et al., 2020)**, overexpression of *IbPrx17* improves salt and drought tolerance in sweet potato **(Zhang et al., 2022),** and overexpression of *OsPrx114* enhances drought tolerance in rice **(Zheng et al., 2023)**. In biotic stress resistance, downregulation of *AtPRX33/AtPRX34* blocks the oxidative burst and causes enhanced susceptibility to pathogens in Arabidopsis **(Daudi et al., 2012)**. Silencing of *CaPO2* (*CaPrx2*) increases disease susceptibility in pepper **(Choi et al., 2007)**. However, it was also shown that suppression of *Ep5C* (*LePrx06*) enhances resistance to the bacterial speck by increasing H_2_O_2_ accumulation in tomatoes **(Coego et al., 2005)**, and overexpression of *OsPrx30* enhances the susceptibility to bacterial blight due to excessive scavenging of H_2_O_2_ **(Liu et al., 2021b)**. These results suggest that Class III Prxs play different roles in biotic stress, catalyzing H_2_O_2_ production as well as catalyzing H_2_O_2_ decomposition. In growth, AtPrx62/AtPrx69 promote root hair growth at low temperature **(Pacheco et al., 2022)**, AtPrx17 participates in lignified tissue formation **(Cosio et al., 2017)**, and AtPrx9/AtPrx40 contributes to tapetal cell wall integrity during development in Arabidopsis **(Jacobowitz et al., 2019)**. Although these reports all indicate that different Class III Prxs function in distinct manners, the biological functions of most Class III Prxs remain unclear. However, one point is clear: Class III Prxs can regulate ROS homeostasis and exhibit a diverse array of functions.

The phytohormone abscisic acid (ABA), accumulated in plants exposed to unfavorable conditions, is the central regulator of abiotic stress tolerance in plants. It coordinates various cellular transduction pathways, enabling plants to activate their defense systems **(Chen et al., 2020; Hewage et al., 2020; Zhu, 2016)**. Among them, the alteration of enzyme activities through post-translational modifications such as phosphorylation, ubiquitination, acetylation, persulfidation, S-nitrosylation, and S-glutathionylation to regulate ROS production and scavenging in response to abiotic or biotic stresses has been widely reported **(Zhang et al., 2022b)**. Among the post-translational modifications, protein phosphorylation plays a crucial role in this process **(Castro et al., 2021; Han et al., 2019)**. calcium (Ca^2+^)/calmodulin (CaM)-dependent protein kinase (CCaMK) decodes the Ca^2+^-centered second messenger system through phosphorylation, which is vital for biotic stress defense and ABA-mediated abiotic stress defense as well as participates in various biological processes, including seed germination and root growth **(Jiang and Ding, 2023; Shi et al., 2014; Shi et al., 2012; Wang et al., 2015)**. In rice, OsDMI3 is a single CCaMK currently reported that is activated by ABA under stress, enhancing downstream antioxidant protection. On the one hand, activated-OsDMI3 mediates the phosphorylation of OsRbohB to generate H_2_O_2_ further and amplify ROS signaling **(Wang et al., 2023)**. On the other hand, activated-OsDMI3 activates the OsMKK1-OsMPK1 cascade pathway by phosphorylating OsMKK1, which transmits signals downstream to scavenge the excess ROS by activating the antioxidant defense system under stress **(Chen et al., 2021)**. Moreover, activated-OsDMI3 indirectly promotes the interaction between OsUXS3 and OsCATB to scavenge H_2_O_2_ by mediating the phosphorylation of OsUXS3, enhancing the CAT activity and thereby scavenging H_2_O_2_ to improve oxidative stress tolerance in rice **(Ni et al., 2022)**. However, redox is a dynamic equilibrium process. OsDMI3 phosphorylates OsRbohB to generate H_2_O_2_ directly, but it phosphorylates OsMKK1 or OsUXS3 to scavenge H_2_O_2_ is an indirect pathway. It is still unclear whether there is a direct Ca^2+^-mediated pathway to scavenge ROS, thus balancing intracellular ROS homeostasis.

Here, we demonstrated that OsPrx20, a member of plant Class III Prxs, is also a phosphorylated substrate of OsDMI3 in ABA signaling. Overexpression of *OsPrx20* directly scavenged ROS to enhance osmotic stress tolerance but reduced the resistance to rice blast in rice. In contrast, knockout of *OsPrx20* was the opposite, indicating that OsPrx20 is a positive regulator of osmotic stress tolerance and a negative regulator of rice blast resistance in rice. Meanwhile, overexpression of *OsPrx20* enlarged spike and grain size, whereas lack of *OsPrx20* could achieve a dwarfism phenotype and smaller spike size. Previous genetic approaches have achieved stress resistance in plants by amplifying the response to stress signals, but this has often come at the cost of inhibiting growth and reducing yield **(Zhang et al., 2020)**. For example, ABA produced under stress conditions enhanced response to abiotic stress, but ABA also inhibits growth and promotes seed dormancy to adapt and survive environmental stresses **(Julkowska and Testerink, 2015; Zhao et al., 2016)**. Remarkably, this limitation is broken in OsPrx20, which was phosphorylated by OsDMI3 to reduce the sensitivity of ABA to seed germination and root growth and to enhance osmotic stress tolerance without affecting spike and grain development, making it a good candidate gene. However, its application is limited because *OsPrx20* overexpression also leads to disease susceptibility. Despite all this, the functional diversity of OsPrx20 and a molecular link between the synchronized enhancement of stress defense and growth provide an exciting research direction for the future.

## 2. Results

### 2.1 OsPrx20 is a target protein of OsDMI3

OsPrx20, a rice Class III Prx, was isolated by screening the rice complementary DNA (cDNA) library using OsDMI3 as a bait. We first verified the authenticity of the interaction between OsPrx20 and OsDMI3 by multiple assays. Yeast two-hybrid (Y2H) assay showed that OsPrx20-AD/OsDMI3-BD, as well as positive control transformed yeast cells grew strongly on SD/-Trp-Leu-His-Ade/+X-ɑ-gal medium. In contrast, the control combinations, OsPrx20-AD/BD-empty or AD-empty/OsDMI3-BD, showed no growth as well as the negative control **(Fig. 1A)**. Glutathione S-transferase (GST) pull-down assay also showed that OsDMI3-GST directly interacted with OsPrx20-His in vitro **(Fig. 1B)**. Luciferase complementation imaging (LCI) assay showed that transiently infiltrated *Nicotiana benthamiana* leaves with constructs encoding the protein fusions OsPrx20-nLUC and OsDMI3-cLUC could observe the reconstitution of firefly luciferase in leaves as well as the positive control. In contrast, no fluorescence was observed when OsPrx20-nLUC/cLUC-vector or nLUC-vector/OsDMI3-cLUC was transferred into the leaf **(Fig. 1C)**. Moreover, bimolecular fluorescence complementation (BiFC) assay in rice protoplasts also showed that OsPrx20 could interact with OsDMI3 to produce yellow fluorescence protein (YFP) in the cytoplasm without overlapping with the blue fluorescence of the DAPI (4’,6-diamidino-2-phenylindole)-stained as a nucleus marker or the FM4-64 (N-(3-triethylammoniopropyl)-4-(6-(4-(diethylamino) phenyl) hexatrienyl) pyridinium dibromide)-stained as a plasma membrane marker, further supporting the fact of the interaction between OsPrx20 and OsDMI3 in vivo **(Fig. 1D)**. The protein expression levels of OsPrx20 and OsDMI3 in tobacco leaves and rice protoplasts were detected by using anti-OsPrx20 and anti-OsDMI3 antibodies in the LCI and BiFC assays **(Fig. S2)**.

**Figure 1.**
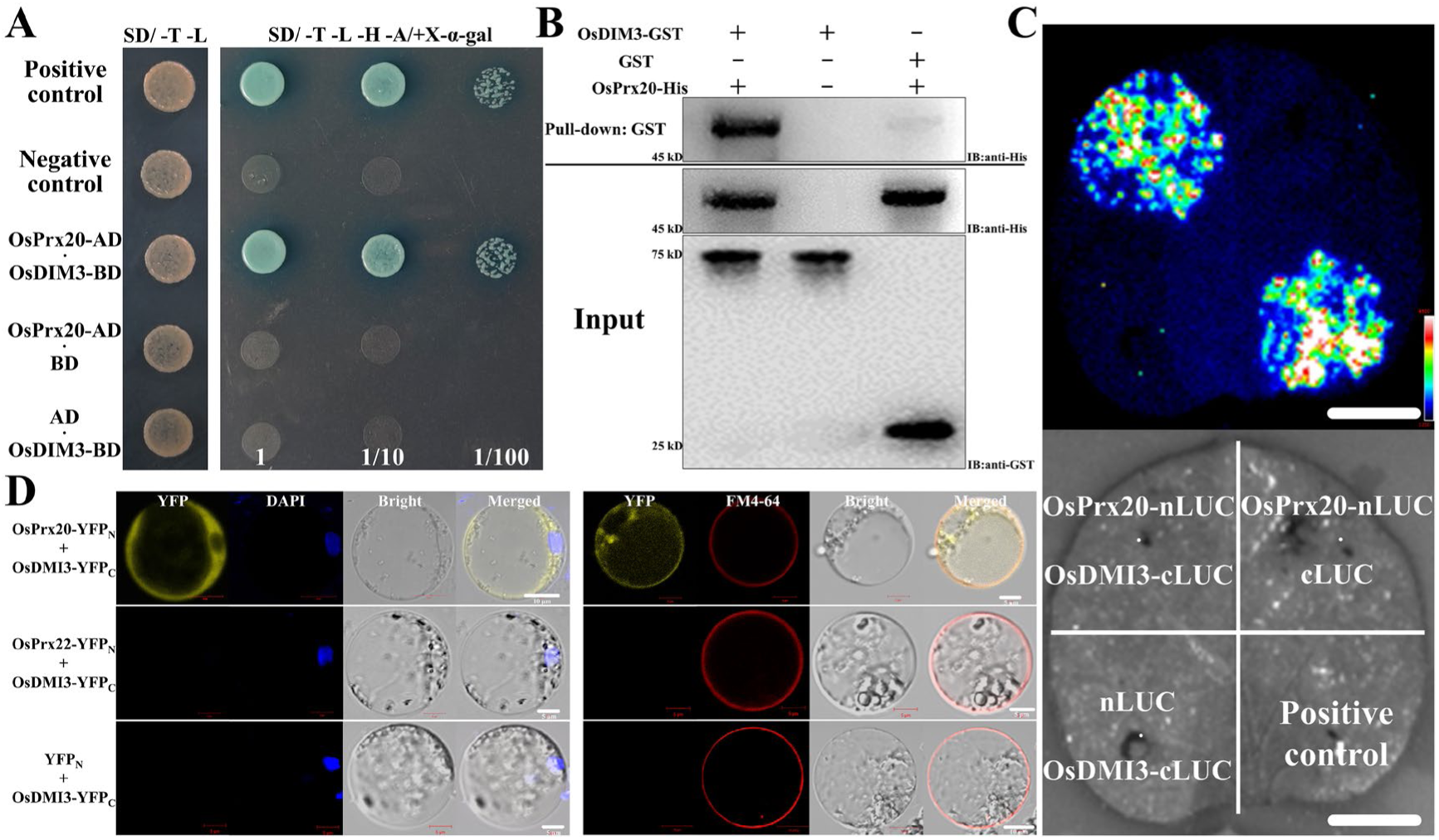
OsPrx20 interacts with OsDMI3. **(A)** Y2H assays for the interaction between OsDMI3 and OsPrx20. The recombinant yeast cells were cultured on SD/-Trp-Leu or SD/-Trp-Leu-His-Ade/+X-ɑ-gal medium and were used for testing the veracity of the interaction between OsDMI3 and OsPrx20. AD (pGADT7) and BD (pGBKT7) are the prey and bait vectors, respectively. The combination of BD-P53/AD-SV40 or BD-Lam/AD-SV40 was used as a positive or negative control. **(B)** GST pull-down assay. The equal amount of OsDMI3-GST or GST was incubated with OsPrx20-His in GST beads. OsPrx20-His was detected with an anti-His antibody, and OsDMI3-GST and GST were detected with an anti-GST antibody. Molecular mass markers in kD were shown on the left. **(C)** LCI assays for the interaction between OsDMI3 and OsPrx20. The tobacco leaves were co-transformed with the constructs OsPrx20-nLUC/OsDMI3-cLUC, and luciferase signals were captured using the Tanon-5200 image system after 3 d *Agrobacterium* infiltration. Combinations of OsPP45-nLUC/OsDMI3-cLUC were used as a positive control, and combinations of OsPrx20-nLUC/Vector-cLUC or Vector-nLUC/OsDMI3-cLUC were used as a negative control. Scale bars, 2 cm. **(D)** BiFC assays for the interaction between OsDMI3 and OsPrx20. The constructs OsPrx20-YFP_N_ and OsDMI3-YFP_C_ were co-transfected into rice protoplasts. Combinations of OsPrx22-YFP_N_/OsDMI3-YFP_C_ or Vector-YFP_N_/OsDMI3-YFP_C_ were used as negative controls. The YFP signals were visualized under ZEISS laser scanning microscopy after incubating for 12 h. The DAPI as a nucleus marker is a dark field showing blue fluorescence, and the FM4-64 as a plasma membrane marker shows red fluorescence. Scale bars, 5/10 μm. The results of all experiments were repeated at least three times with similar results.

What and where is the role of OsPrx20 in rice? We analyzed the transcript levels of *OsPrx20* in different tissues of rice and found that it was expressed in leaves, stems, and roots, with the highest transcript levels in stems and almost none in the elongation and meristematic zones of roots **(Fig. S1A)**. GUS staining results in the transgenic plants of *proOsPrx20*-GUS were consistent with the RT-qPCR results, with the deepest GUS staining in the stems, followed by the leaves and maturation zone of roots. In contrast, the elongation zone and meristematic zone of roots were not stained **(Fig. S1B)**. Following this, transient expression of the pXZP008-*OsPrx20*-YFP recombinant construct in rice protoplasts was performed. We found the OsPrx20-YFP was located in the cytoplasm and overlapped with neither the blue fluorescence of DAPI-stained as a nuclear marker nor the red fluorescence of FM4-64-stained as a cell membrane marker **(Fig. S1C)**. In contrast, the YFP was present throughout the cell by transforming the empty vector (pXZP008-YFP) alone. The result was further supported by heterologous expression in tobacco cells, where OsPrx20 was only localized in the cytoplasm and overlapped with neither DAPI-stained nor FM4-64-stained **(Fig. S1D)**.

### 2.2 OsPrx20 positively regulates osmotic stress tolerance but negatively regulates blast resistance in rice

Previous studies have been reported for CCaMKs and Class III Prxs in response to abiotic and biotic stresses **(Chen et al., 2021; Liu et al., 2021b; Ni et al., 2019; Wang et al., 2015; Wang et al., 2023; Zhang et al., 2022; Zheng et al., 2023)**. To determine whether OsPrx20 also has a regulatory effect on abiotic and biotic stress, two knockout mutant lines and two overexpression transgenic lines of *OsPrx20* (*osprx20* KO1, *osprx20* KO2, *OsPrx20* OE1, and *OsPrx20* OE2) were generated **(Fig. S3A and S3B)** and these mutants and transgenic lines were treated in 20% polyethylene glycol (PEG)-6000 solution to simulate abiotic stress and infected with *M. oryzae* strain Guy11 to simulate biotic stress. The results of abiotic stress showed a lower osmotic stress tolerance in *osprx20*-KO after treatment in PEG solution for 7 d compared with WT, while a higher tolerance in *OsPrx20*-OE **(Fig. 2A)**. After being allowed to recover by re-watering for a week, *osprx20*-KO had a significantly lower survival rate than WT, while *OsPrx20*-OE was higher than WT, the survival rate of *osprx20*-KO was only 20-30%, much lower than the 50%-55% of WT and the 75-80% of *OsPrx20*-OE **(Fig. 2B).** Osmotic stress caused excessive accumulation of ROS in plant cells, resulting in oxidative damage. Here, we found higher H_2_O_2_ levels and lower total peroxidase (POD) activity in *osprx20*-KO after PEG treatment than in WT. Conversely, *OsPrx20*-OE had lower H_2_O_2_ levels and higher POD activity than WT **(Fig. S4A and S4B)**. Meanwhile, the accumulation of ROS in the leaves of *osprx20*-KO, *OsPrx20*-OE, and WT exposed to PEG solution was observed using 3,3-diaminobenzidine (DAB) staining. The results showed no DAB staining in the leaves of *osprx20*-KO, *OsPrx20*-OE, and WT without treatment. Following PEG treatment, lighter DAB staining in *osprx20*-KO and darker DAB staining in *OsPrx20*-OE than WT were observed **(Fig. S4C)**. Malondialdehyde (MDA) content and electrolyte leakage rate, two critical indicators of oxidative damage, were also measured here. These results showed that there was a significantly higher MDA content and electrolyte leakage rate in *osprx20*-KO than that in WT and *OsPrx20*-OE after PEG treatment, while *OsPrx20*-OE had the lowest **(Fig. S4D and S4E)**. Taken together, these results indicated that lack of *OsPrx20* significantly increased ROS accumulation under osmotic stress, whereas overexpression of *OsPrx20* significantly alleviated oxidative damage. However, the biotic stress results showed that *osprx20*-KO was significantly more resistant to rice blast infection than WT, whereas *OsPrx20*-OE was more sensitive. After the rice leaves were spray-inoculated with Guy11 spores for 7 d, there was the lowest level of lesions on *osprx20*-KO leaves with the lesion type concentrated in type 1, whereas *OsPrx20*-OE leaves with the lesion type concentrated in type 3 and type 4 exhibited more level of lesions than both WT with the lesion type concentrated in type 2 and *osprx20*-KO **(Fig. 2C and 2D)**. The propagation of *M. oryzae* on the infected leaves was quantified by qRT-PCR using primers specific to *MoPot2*, an *M. oryzae* housekeeping gene. In agreement with the resistant phenotype, the least hyphae propagation was detected in leaves of *osprx20*-KO than in WT and *OsPrx20*-OE **(Fig. 2E)**. Meanwhile, the expression of *OsPrx20* and the levels of OsPrx20 protein showed significant down-regulation under rice blast infection **(Fig. 2F and 2G)**. These results obtained from abiotic and biotic stresses indicate that OsPrx20 is a positive regulator of osmotic stress tolerance but a negative regulator of rice blast immunity.

**Figure 2.**
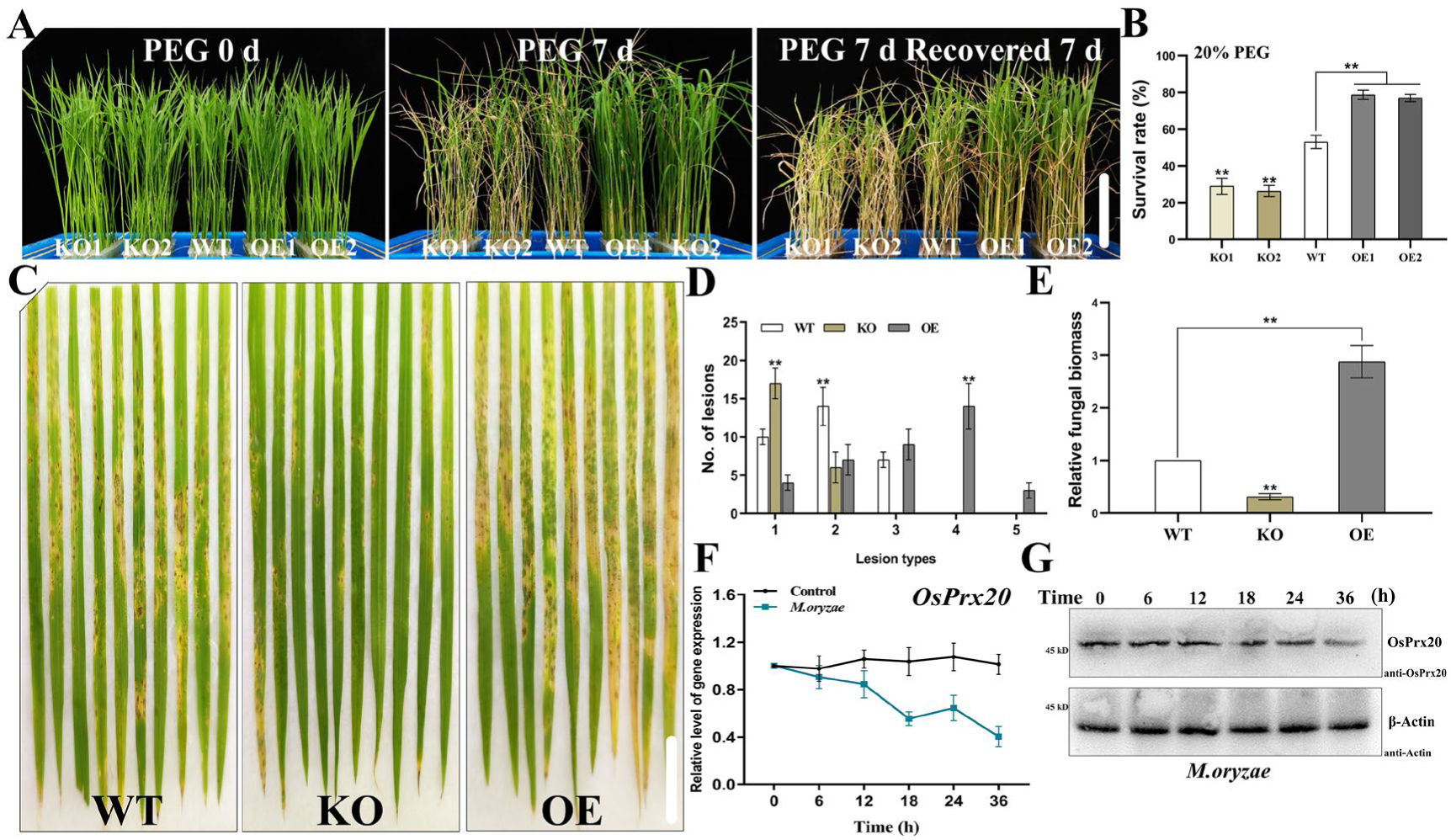
Phenotypic analyses of *osprx20-KO*, *OsPrx20*-OE, and WT exposed to osmotic or rice blast stress. **(A)** Phenotypes of *osprx20-*KO, *OsPrx20*-OE, and WT exposed to osmotic stress. 15-day-old rice seedlings were treated with 20% PEG-6000 solution for 7 d and then recovered by re-watering for 7 d. Scale bars, 10 cm. **(B)** The survival rate (%) of *osprx20-*KO, *OsPrx20*-OE, and WT after recovery by re-watering for 7 d is shown in (A). **(C)** Phenotypes of *osprx20-*KO, *OsPrx20*-OE, and WT leaves infected with rice blast. 15-day-old rice seedlings were conidial suspensions (1 × 10^5^ spores/mL) of the Guy11 strain, and photographs were taken after 7 days post-inoculation. Scale bar, 2 cm. **(D)** Lesion types were counted (Type 1, uniformly dark brown pinpoint lesions without visible centers; Type 2, small lesions with distinct tan centers surrounded by a darker brown margin; Type 3, small eyespot lesions approximately 1 mm in length with tan centers surrounded by dark brown margins; Type 4, intermediate size eyespot lesions, about 2–3 mm in length; Type 5, large eyespot lesions that attain the maximum size seen). Lesion numbers within an area of 4 cm^2^ were counted. **(E)** Relative fungal biomass is determined by examining the expression level of the *MoPot2* against the *OsUbiquitin* DNA level. The expression level of *MoPot2* in WT was defined as 1.0. **(F)** Expression of *OsPrx20* in the leaves of WT infected with rice blast fungus. 15-day-old rice seedlings were infected with Guy11 at different times. Expression levels were defined in treatment 0 min as 1.0. The *OsGAPDH* was selected as an internal control for normalization. **(G)** Expression of OsPrx20 in the leaves of WT infected with rice blast fungus. Total proteins extracted from WT leaves were detected by IB with an anti-OsPrx20 antibody after being infected with Guy11 at different times. β-Actin was used as total protein internal control. Molecular mass markers in kD were shown on the left. The results of all experiments were repeated at least three times with similar results. Approximately 48 seedlings of each rice material were used per replicate in (A, C). Data in (B, D, E, and F) are means ± SD (n = 3). Significant differences are indicated: **, *P* < 0.01 (one-way ANOVA and Tukey’s multiple comparison tests).

### 2.3 Overexpression of *OsPrx20* enlarges spike and grain size, whereas lack of *OsPrx20* leads to dwarfism and a smaller spike

Plant height and tiller number are critical agronomic traits for measuring rice growth and harvest, directly affecting the spikes and yield. At rice maturity, we found that *osprx20*-KO exhibited significant dwarfism compared with WT, whereas *OsPrx20*-OE showed no significant discrepancy in plant height compared with WT, suggesting that OsPrx20 is required for stem elongation and also indirectly explaining the expression of *OsPrx20* mainly in the stems **(Fig. 3A and 3B)**. However, neither lack nor overexpression of *OsPrx20* caused a change in tiller number **(Fig. 3C)**. Lignin participates in the growth-defense trade-offs, while Class III Prxs can promote lignin biosynthesis **(Almagro et al., 2009; Shigeto and Tsutsumi, 2016; Xie et al., 2018)**. To investigate a potential connection between OsPrx20 and lignin biosynthesis, the lignin content in leaves, stems, and roots of 15-day-old and 10-week-old *osprx20*-KO, *OsPrx20*-OE, and WT plants were determined. There was no significant difference in the content of lignin in leaves, stems, and roots of the 15-day-old seedlings **(Fig. S5A)**. However, lack of *OsPxr20* decreased lignin content in adult stems and roots, whereas overexpression of *OsPxr20* increased lignin content in adult stems and roots **(Fig. S5B)**. But OsPrx20 localized in the cytoplasm **(Fig. S3C)**, suggesting that OsPrx20 might indirectly modulate lignification in the stems and roots during rice growth. Although the lack of *OsPrx20* could lead to a dwarfism phenotype, the reduction in lignification degree was unlikely to improve the ability to resist lodging.

**Figure 3.**
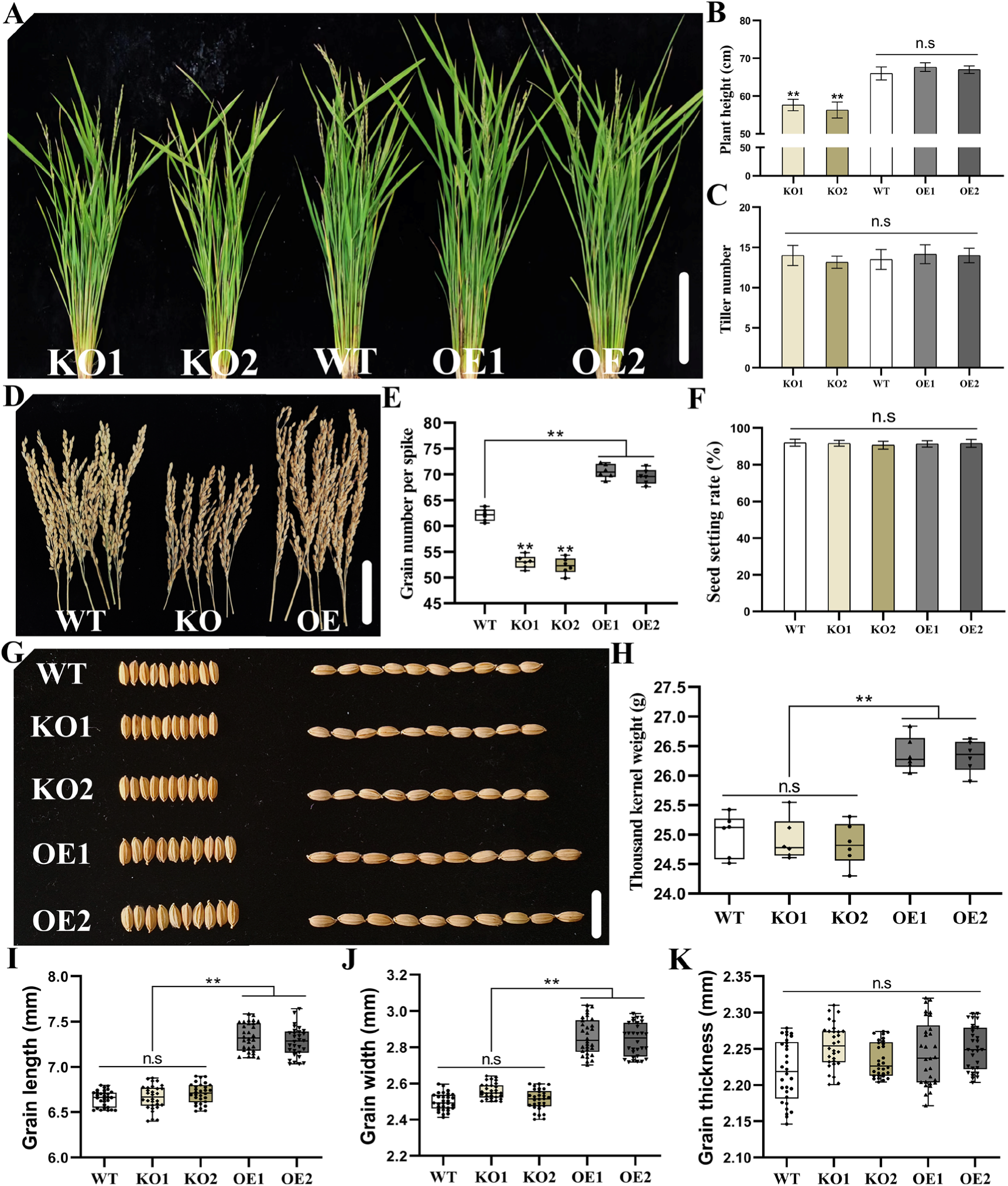
Phenotypic analyses of *osprx20*-KO, *OsPrx20*-OE, and WT in plant, spike, and grain. **(A)** Plant height phenotype of *osprx20-*KO, *OsPrx20*-OE, and WT were photographed at 10 weeks old. Scale bar, 20 cm. **(B)** Plant height. **(C)** Tiller number. **(D)** Single-spike phenotype of *osprx20*-KO, *OsPrx20*-OE, and WT. **(E)** Grain number per spike. **(F)** Seed setting rate per spike. **(G)** Grain phenotype of *osprx20-*KO, *OsPrx20*-OE, and WT. **(H)** Thousand kernel weights. **(I)** Grain length. **(J)** Grain width. **(K)** Grain thickness. The results of all experiments were repeated at least three times. Data in (B, C, E, F, and H) are means ± SD (*n* = 6). Data in (I, J, and K) are means ± SD (*n* = 30). Significant differences are indicated: **, *P* < 0.01; no significant differences are indicated: n.s (one-way ANOVA and Tukey’s multiple comparison tests).

Others important agronomic traits caused by *OsPrx20* was revealed by comparing the spike and grain among *osprx20*-KO, *OsPrx20*-OE, and WT. The size of *osprx20*-KO spikes was significantly smaller than WT, whereas *OsPrx20*-OE was significantly larger **(Fig. 3D)**. Although knockout or overexpression of *OsPrx20* did not cause the difference in seed setting rate, it directly affected the grain number per spike with a significantly lower in *osprx20*-KO than in WT, whereas a higher in *OsPrx20*-OE **(Fig. 3E and 3F).** The length, width, and thicknesses between *osprx20*-KO and WT seeds were insignificant. The thicknesses also had no significance between *OsPrx20*-OE and WT seeds, but the length and width of *OsPrx20*-OE seeds increased by ∼9% and ∼12% compared with the WT **(Fig. 3I, 3J, and 3K)**. These directly led to the discrepancy of the thousand kernel weight among *osprx20*-KO, *OsPrx20*-OE, and WT seeds, with a weightier in *OsPrx20*-OE seeds than in WT and *osprx20*-KO **(Fig. 3H)**. Although there was no significance in the thousand kernel weight between *osprx20*-KO and WT, it still resulted in a decline in single plant yield due to its reduced grain number per spike. Taken together, these results indicated that OsPrx20 is a positive regulator of growth and grain development. In natural conditions, lack of *OsPrx20* caused dwarfism and a smaller spike in rice, whereas overexpression of *OsPrx20* enlarged spike size and grain fullness without affecting plant height and tiller number.

### 2.4 OsPrx20 Thr-244 phosphorylation is critical for osmotic stress tolerance and for rice blast resistance

Previous studies have shown that OsDMI3 can phosphorylate multiple substrates, including OsMKK1, OsUXS3, and OsRbohB **(Chen et al., 2021; Ni et al., 2022; Wang et al., 2023)**. To determine whether OsPrx20 is a phosphorylated substrate of OsDMI3, recombinant OsPrx20-His fragment was incubated with OsDMI3-GST. Polypeptide fragments of OsPrx20 were determined by liquid chromatography-tandem mass spectrometry (LC-MS/MS) analysis. Thr-244 in OsPrx20 was identified as the site phosphorylated by OsDMI3 in vitro **(Fig. 4A)**. Further, Thr-244 in OsPrx20 was substituted with alanine (Ala) to make it non-phosphorylatable and in vitro kinase assays showed that OsPrx20^T244A^-His was not phosphorylated by OsDMI3-GST **(Fig. 4B)**. These results indicated that Thr-244 of OsPrx20 is the phosphorylation site for OsDMI3 in vitro. Then, a direct analysis of the function of OsPrx20 phosphorylation at Thr-244 site using genetic materials was performed. The stable transgenic rice lines overexpressing *OsPrx20^T244A^*-OE (Thr-244 of OsPrx20 was mutated to alanine (Ala) to create non-phosphorylated overexpression transgenic rice, A1 and A2) or *OsPrx20^T244D^*-OE (Thr-244 of OsPrx20 was mutated to aspartate (Asp) to create phosphomimetic overexpression transgenic rice; D1 and D2) were generated in the WT background. In these transgenic lines, the expression levels of *OsPrx20* were similar **(Fig. S3B)**. These transgenic lines, as well as *osprx20*-KO lines, *OsPrx20*-OE lines, and WT plants, were treated with PEG, or *M. oryzae*. Although *OsPrx20*-OE, *OsPrx20^T244A^*-OE, and *OsPrx20^T244D^*-OE were all better in phenotype and survival rate after treatment in PEG solution for a week then being allowed to recover by re-watering for a week compared with WT, a significantly better osmotic stress tolerance was observed in *OsPrx20^T244D^*-OE compared with *OsPrx20*-OE, whereas a weaker in *OsPrx20^T244A^*-OE **(Fig. 4C)**. *OsPrx20^T244D^*-OE had the highest survival rate of ∼85%, far higher than the ∼75% of *OsPrx20*-OE and the ∼60% of *OsPrx20^T244A^*-OE **(Fig. 4D).** The measurements of oxidative damage indicators were also consistent with the phenotypes. After treatment with PEG, *OsPrx20^T244D^*-OE possessed the lowest MDA content, the lowest electrolyte leakage rate, and the weakest DAB staining, whereas compared with *OsPrx20*-OE, *OsPrx20^T244A^*-OE exhibited stronger oxidative damage and DAB staining **(Fig. S6)**. Next, we further assayed the activity of OsPrx20 using a POD assay kit. Without treatment, *OsPrx20^T244D^*-OE already had a higher OsPrx20 activity compared with other materials. After treatment, the highest level of OsPrx20 activity also in *OsPrx20^T244D^*-OE after treatment compared with the other materials, but a lower level of OsPrx20 activity in *OsPrx20^T244A^*-OE compared with *OsPrx20*-OE **(Fig. 4E)**. Interestingly, in vitro prokaryotic expression of OsPrx20, neither OsPrx20, OsPrx20^T244A^, nor OsPrx20^T244D^ had POD activity **(Fig. S7)**. This suggested that OsPrx20 probably requires post-translational modification in vivo to possess peroxidase activity. Together, these results indicated that phosphorylation of OsPrx20 at Thr-244 site activated the ability of OsPrx20 as a peroxidase to scavenge ROS, thereby reducing the accumulation of ROS under osmotic stress. However, phosphorylation of OsPrx20 at Thr-244 caused a further increase in susceptibility to rice blast. Although *OsPrx20-OE*, *OsPrx20^T244A^*-OE, and *OsPrx20^T244D^*-OE all exhibited more lesions than WT and *osprx20*-KO after spray-inoculating with Guy11 spores, *OsPrx20^T244D^*-OE had the most lesions with lesion type developed from type 4 to type 5 and the most accumulation of hyphae, whereas *OsPrx20^T244A^*-OE was mitigated with the lesion type concentrated in type 3 compared with *OsPrx20*-OE **(Fig. 4F, 4G, and 4H)**. Meanwhile, The content of H_2_O_2_ in the leaves showed that *OsPrx20^T244D^*-OE leaves had the lowest levels of H_2_O_2_ even after infection, while *osprx20*-KO leaves had the highest **(Fig. S8A)**. It was also confirmed by DAB staining, which showed the lightest DAB staining in the most severely infected *OsPrx20^T244D^*-OE leaves, whereas the deepest in *osprx20*-KO leaves **(Fig. S8B)**. The most likely of this phenomenon was excess ROS scavenging caused by the increased POD activity following the phosphorylation of Thr-244 in OsPrx20, which nicely explained why the expression of OsPrx20 was down-regulated under rice blast infection **(Fig. 2H and 2I)**. Further comparing the spike and grain among *OsPrx20*-OE, *OsPrx20^T244A^*-OE, and *OsPrx20^T244D^*-OE, we found that either spikes or seeds had no significant among them. These also directly led to the thousand kernel weight having no change among them **(Fig. S9)**. Similarly, the lignin content did not vary in *OsPrx20^T244A^*-OE and *OsPrx20^T244D^*-OE compared to *OsPrx20*-OE **(Fig. S10)**. Taken together, overexpression of *OsPrx20* contributed to rice grain development in natural conditions, but phosphorylation of OsPrx20 at the Thr-244 site, whether or not, did not further enhance this effect in rice. These also indicated that OsPrx20 Thr-244 phosphorylation is a specific phosphorylation site under osmotic stress.

**Figure 4.**
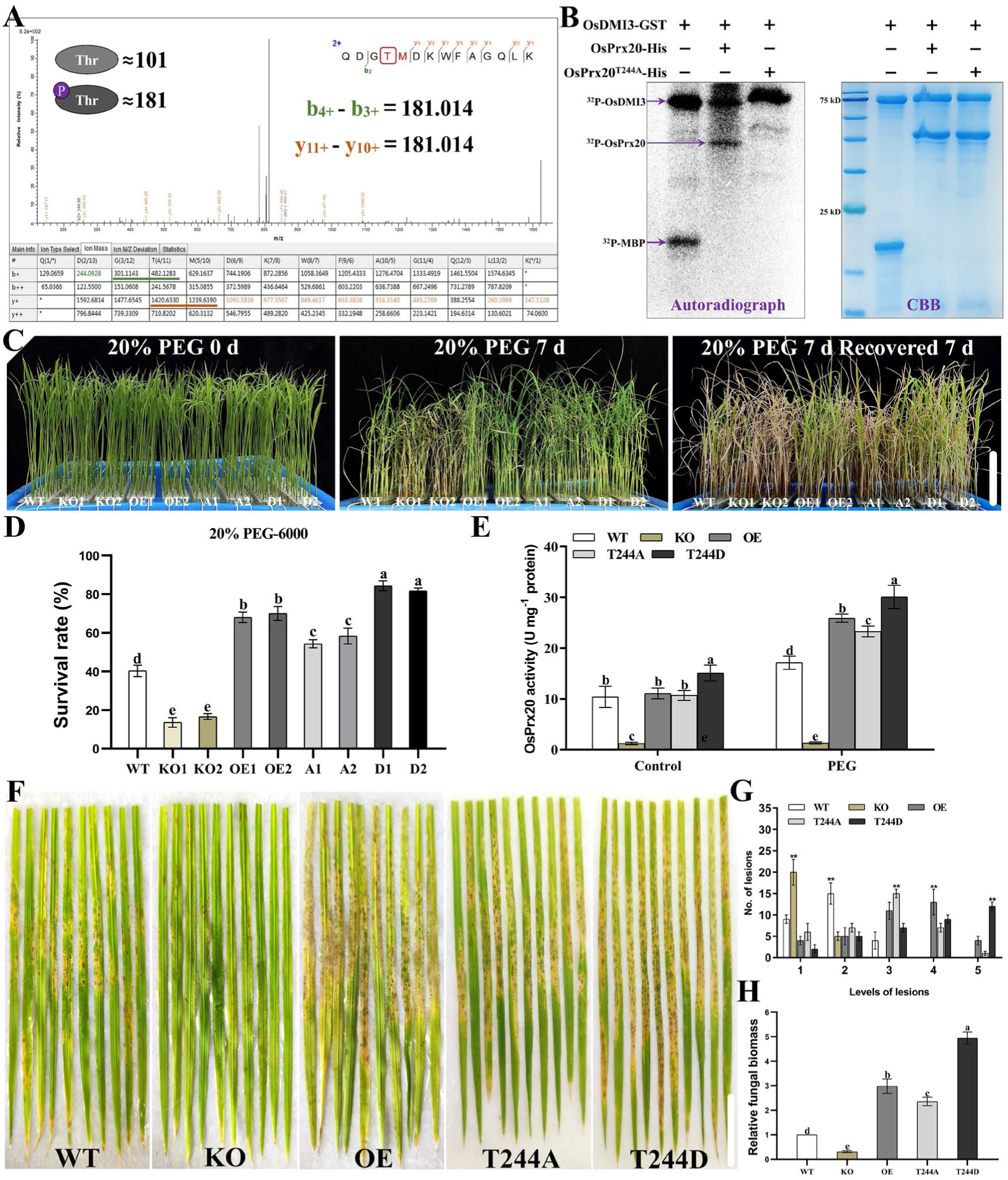
Function of OsPrx20 Thr-244 phosphorylation in rice exposed to osmotic or rice blast stress. **(A)** Identifies the sites of OsPrx20 phosphorylated by OsDMI3 in vitro using Liquid chromatography-tandem mass spectrometry (LC-MS/MS). The purified OsDMI3-GST and OsPrx20-His proteins were incubated in a kinase reaction buffer, and Thr-244 in OsPrx20 was identified as the phosphorylated site. **(B)** Phosphorylation of OsPrx20 and mutant OsPrx20 (T244A) by OsDMI3 was analyzed in vitro. OsPrx20-His, OsPrx20^T244A^-His, and OsDMI3-GST proteins were expressed in *Escherichia coli.* OsDMI3-GST as a kinase, OsPrx20-His, and mutated OsPrx20^T244A^-His proteins were used as substrates and subjected to an in-gel kinase assay. Myelin basic protein (MBP) was used as a positive substrate. The gel staining by using the Coomassie Brilliant Blue. **(C)** Phenotypes of *osprx20*-KO, *OsPrx20*-OE, *OsPrx20^T244A^*-OE, *OsPrx20^T244D^*-OE, and WT exposed to osmotic stress. 15-day-old rice seedlings were treated in 20% PEG-6000 solution for 7 d and then recovered by re-watering for 7 d. Scale bars, 10 cm. **(D)** The survival rate (%) of *osprx20-*KO, *OsPrx20*-OE, and WT after recovery by re-watering for 7 d is shown in (C). **(E)** The OsPrx20 activity in *osprx20*-KO, *OsPrx20*-OE, *OsPrx20^T244A^*-OE, *OsPrx20^T244D^*-OE, and WT after treated in 20% PEG-6000 solution for 2 d. Assay OsPrx20 activity using the Amplex® Red Hydrogen Peroxide/Peroxidase Assay Kit. The peroxidase standard curve was made with the Horseradish peroxidase (HRP) standard. **(F)** Phenotypes of *osprx20-KO*, *OsPrx20*-OE, *OsPrx20^T244A^*-OE, *OsPrx20^T244D^*-OE, and WT leaves infected with rice blast. 15-day-old rice seedlings were conidial suspensions (1 × 10^5^ spores/mL) of the Guy11 strain, and photographs were taken after 7 days post-inoculation. Scale bar, 2 cm. **(G)** Lesion types were counted (Type 1, uniformly dark brown pinpoint lesions without visible centers; Type 2, small lesions with distinct tan centers surrounded by a darker brown margin; Type 3, small eyespot lesions approximately 1 mm in length with tan centers surrounded by dark brown margins; Type 4, intermediate size eyespot lesions, about 2–3 mm in length; Type 5, large eyespot lesions that attain the maximum size seen). Lesion numbers within an area of 4 cm^2^ were counted. **(H)** Relative fungal biomass is determined by examining the expression level of the *MoPot2* against the *OsUbiquitin* DNA level. The expression level of *MoPot2* in WT was defined as 1.0. The results of all experiments were repeated at least three times with similar results. Approximately 48 seedlings of each rice material were used per replicate in (C, F). Data in (D, E, G, and H) are means ± SD (n = 3). Significant differences are indicated: different lowercase letters, *P* < 0.05 or **, *P* < 0.01 (one-way ANOVA and Tukey’s multiple comparison tests).

### 2.5 OsPrx20 Thr-244 phosphorylation reduces the sensitivity of ABA to seed germination and root growth

Previous studies have shown that OsDMI3 is an important regulator of ABA signaling **(Chen et al., 2021; Ni et al., 2019; Shi et al., 2014; Wang et al., 2023)**. To determine whether OsPrx20 is also involved in the response to ABA signaling, *osprx20*-KO, *OsPrx20*-OE, and WT seeds were treated with different concentrations of ABA to analyze seed germination. The results showed that the sensitivity of *OsPrx20*-OE to ABA during seed germination was significantly reduced compared with WT, whereas the sensitivity of *osprx20*-KO to ABA was enhanced considerably **(Fig. S11)**, indicating that OsPrx20 negatively regulates ABA response in seed germination. To determine the function of OsPrx20 Thr-244 phosphorylation in ABA signaling, *osprx20*-KO, *OsPrx20*-OE, *OsPrx20^T244A^*-OE, *OsPrx20^T244D^*-OE, and WT seeds were treated with different concentrations of ABA. Under the non-treated condition, there were no obvious differences in seed germination **(Fig. 5A and 5B)** and primary root growth **(Fig. 5C and 5D)** among these materials. After ABA treatment, the ABA sensitivity of seed germination and primary root growth was significantly reduced in *OsPrx20*-OE, *OsPrx20^T244A^*-OE, and *OsPrx20^T244D^*-OE compared with WT. However, when compared with *OsPrx20*-OE, *OsPrx20^T244D^*-OE exhibited a significantly reduced sensitivity to ABA, but *OsPrx20^T244A^*-OE displayed a markedly enhanced sensitivity to ABA, indicating that OsPrx20 Thr-244 phosphorylation can reduce ABA-mediated inhibition of seed germination and primary root growth.

**Figure 5.**
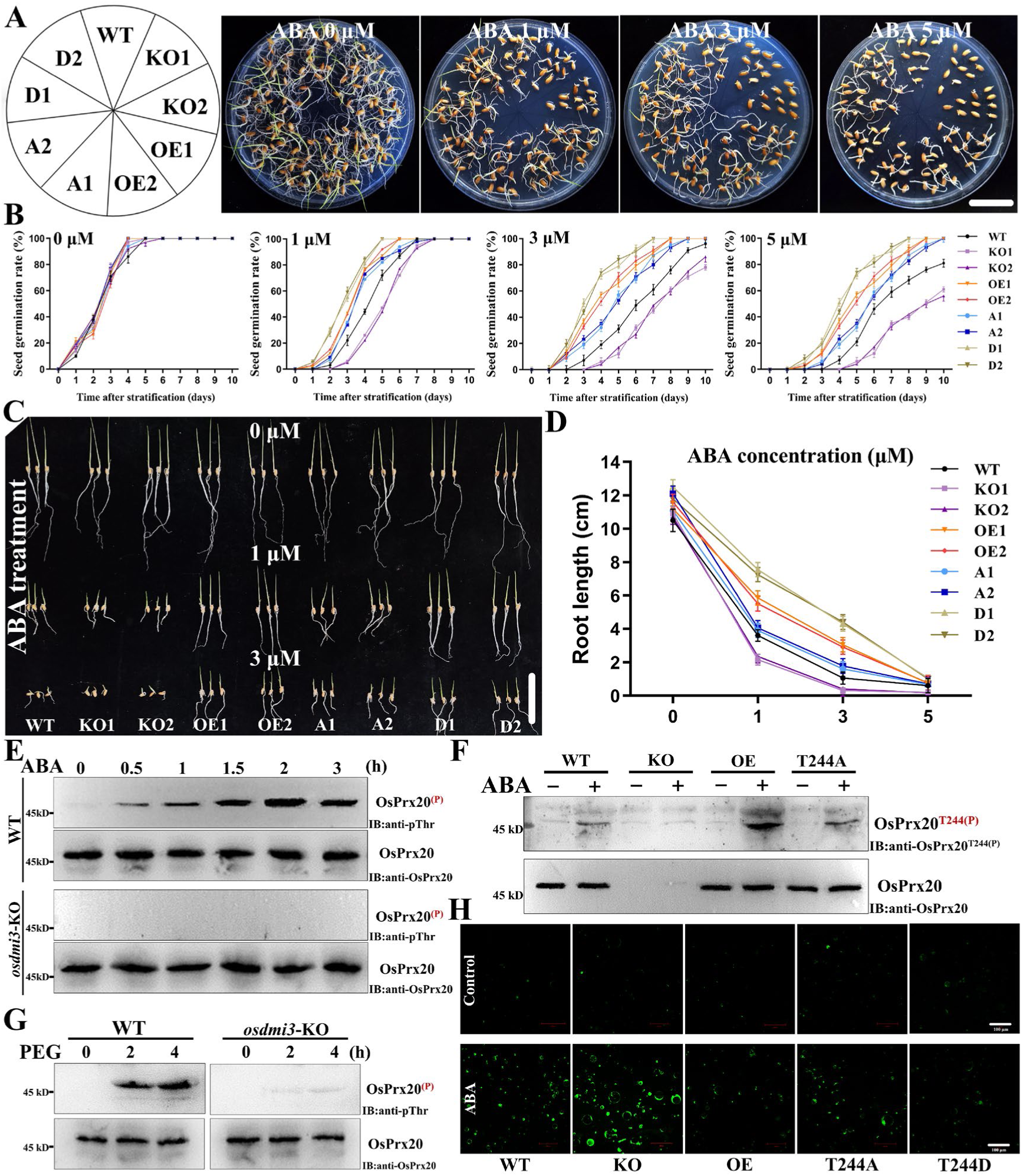
Function of OsPrx20 Thr-244 phosphorylation in ABA signaling. **(A)** Seed germination phenotypes of *osprx20-KO*, *OsPrx20*-OE, *OsPrx20^T244A^*-OE, *OsPrx20^T244D^*-OE, and WT were photographed after treatment with different concentrations of ABA (0, 1, 3, and 5 μM) for 10 d. Scale bar, 3 cm. **(B)** The seed germination rates are shown in (A). **(C)** Primary root growth phenotypes of *osprx20-KO*, *OsPrx20*-OE, *OsPrx20^T244A^*-OE, *OsPrx20^T244D^*-OE, and WT were photographed after treatment with different concentrations of ABA (0, 1, and 3 μM) for 5 d. Scale bar, 5 cm. **(D)** Primary root lengths of the materials shown in (C) grown in different concentrations of ABA as indicated for 10 d. **(E, G)** Phosphorylation levels of OsPrx20 Thr in *vivo* after ABA or PEG treatment. Total proteins were extracted from WT or *osdmi3*-KO plants after being treated in 100 μM ABA or 20% PEG-6000 solution for different times, then immunoprecipitation of OsPrx20 using an anti-OsPrx20 antibody. Phosphorylation of OsPrx20 Thr was tested by IB with an anti-pThr antibody. The input of OsPrx20 was analyzed by IB with an anti-OsOsPrx20 antibody. Molecular mass markers in kD were shown on the left. **(F)** Phosphorylation levels of OsPrx20 Thr-244 in *osprx20*-KO, *OsPrx20*-OE, *OsPrx20^T244A^*-OE, and WT after ABA treatment. Total proteins were extracted from *osprx20*-KO, *OsPrx20*-OE, *OsPrx20^T244A^*-OE, and WT plants after being treated in 1 mM ABA solution for 2 h, then immunoprecipitation of OsPrx20 using an anti-OsPrx20 antibody. Phosphorylation of OsPrx20 Thr-244 was tested by IB with an anti-phospho-OsPrx20^Thr244^ antibody. The input of OsPrx20 was analyzed by IB with an anti-OsPrx20 antibody. Molecular mass markers in kD were shown on the left. **(H)** The fluorescence of ROS was detected in *osprx20*-KO, *OsPrx20*-OE, *OsPrx20^T244A^*-OE, *OsPrx20^T244D^*-OE, and WT rice protoplasts after being treated in 1 μM ABA using CM-H_2_DCFDA. Scale bar, 100 μm. The results of all experiments were repeated at least three times with similar results. Data in (B) are means ± SD (n = 3). Data in (D) are means ± SD (n = 12).

To determine if OsDMI3 is responsible for the phosphorylation of Thr-244 in OsPrx20 in rice, the total protein extracted from WT or *osdmi3*-KO was treated in ABA (100 μM) or 20% PEG-6000 solution for different times, followed by immunoprecipitation of OsPrx20. An anti-phospho-threonine (anti-pThr) antibody was used to detect the phosphorylation of OsPrx20. The IB analysis showed that the phosphorylation of OsPrx20 significantly increased with time and peaked at 2 h during ABA or PEG treatment in WT **(Fig. 5E and 5G)**. However, no phosphorylation bands of OsPrx20 were detected in *osdmi3*-KO, indicating that OsDMI3 is indeed required for ABA/PEG-induced phosphorylation of OsPrx20 at Thr-244 site in rice. Further, a specific anti-phospho-OsPrx20^T244^ antibody was prepared. *osprx20*-KO, *OsPrx20*-OE, *OsPrx20^T244A^*-OE, and WT were treated with ABA, and OsPrx20 Thr-244 phosphorylation was tested by immunoblotting with an anti-phospho-OsPrx20^Thr244^ antibody. Under non-treated condition, OsPrx20 Thr-244 phosphorylation was not detected in all materials. ABA treatment induced an increase in OsPrx20 Thr-244 phosphorylation in *OsPrx20*-OE, *OsPrx20^T244A^*-OE, and WT plants. After ABA treatment for 2 h, the phosphorylation intensity of *OsPrx20^T244A^*-OE was essentially the same as that of WT (*OsPrx20^T244A^*-OE in the background of WT), whereas *OsPrx20*-OE showed a significant enhancement in phosphorylation intensity **(Fig. 5F)**. In contrast, ABA-induced OsPrx20 Thr-244 phosphorylation was completely abolished in *osprx20*-KO. These results indicate that ABA can indeed induce OsPrx20 Thr-244 phosphorylation.

To determine whether ABA-induced OsPrx20 Thr-244 phosphorylation can regulate the level of ROS, CM-H_2_DCFDA (2′,7′-Dichlorodihydrofluorescein diacetate), a ROS-specific fluorescent probe, was used to detect ROS levels in rice protoplasts of *osprx20*-KO, *OsPrx20*-OE, *OsPrx20^T244A^*-OE, *OsPrx20^T244D^*-OE, and WT after treatment with ABA (1 μM). Under non-treated condition, almost no fluorescence of ROS was detected in the protoplasts of all materials **(Fig. 5H)**. After ABA treatment, the fluorescence of ROS in *OsPrx20*-OE, *OsPrx20^T244A^*-OE, and *OsPrx20^T244D^*-OE was attenuated compared with WT, with the weakest fluorescence intensity in *OsPrx20^T244D^*-OE. In contrast, the strongest fluorescence was detected in *osprx20*-KO, suggesting that OsPrx20 Thr-244 phosphorylation can reduce ABA-induced increase in ROS level.

### 2.6 The OsDMI3-OsPrx20 pathway is critical for ABA response and for osmotic stress tolerance

To further confirm that it is indeed the OsDMI3-OsPrx20 pathway that regulates the ABA responses in seed germination and root growth, *osdmi3*-KO, *osdmi3*/*OsPrx20*-OE, *osdmi3*/*OsPrx20^T244D^*-OE (which were generated in the *osdmi3*-KO background; **Fig. S3B**), and WT seeds were treated with different concentrations of ABA, and seed germination and root length of these materials under ABA treatment were analyzed. *osdmi3*-KO, *osdmi3*/*OsPrx20*-OE, and *osdmi3*/*OsPrx20^T244D^*-OE all showed reduced sensitivity to ABA in seed germination **(Fig. 6A and 6B)** and primary root growth **(Fig. 6C and 6D)** compared with WT. Compared with *osdmi3*-KO, *osdmi3/OsPrx20*-OE exhibited a similar sensitivity to ABA, but *osdmi3*/*OsPrx20^T244D^*-OE displayed a greatly reduced sensitivity to ABA. These results indicate that the OsPrx20-mediated ABA responses are dependent on the action of OsDMI3 and OsDMI3-mediated OsPrx20 Thr-244 phosphorylation is important for the ABA responses.

**Figure 6.**
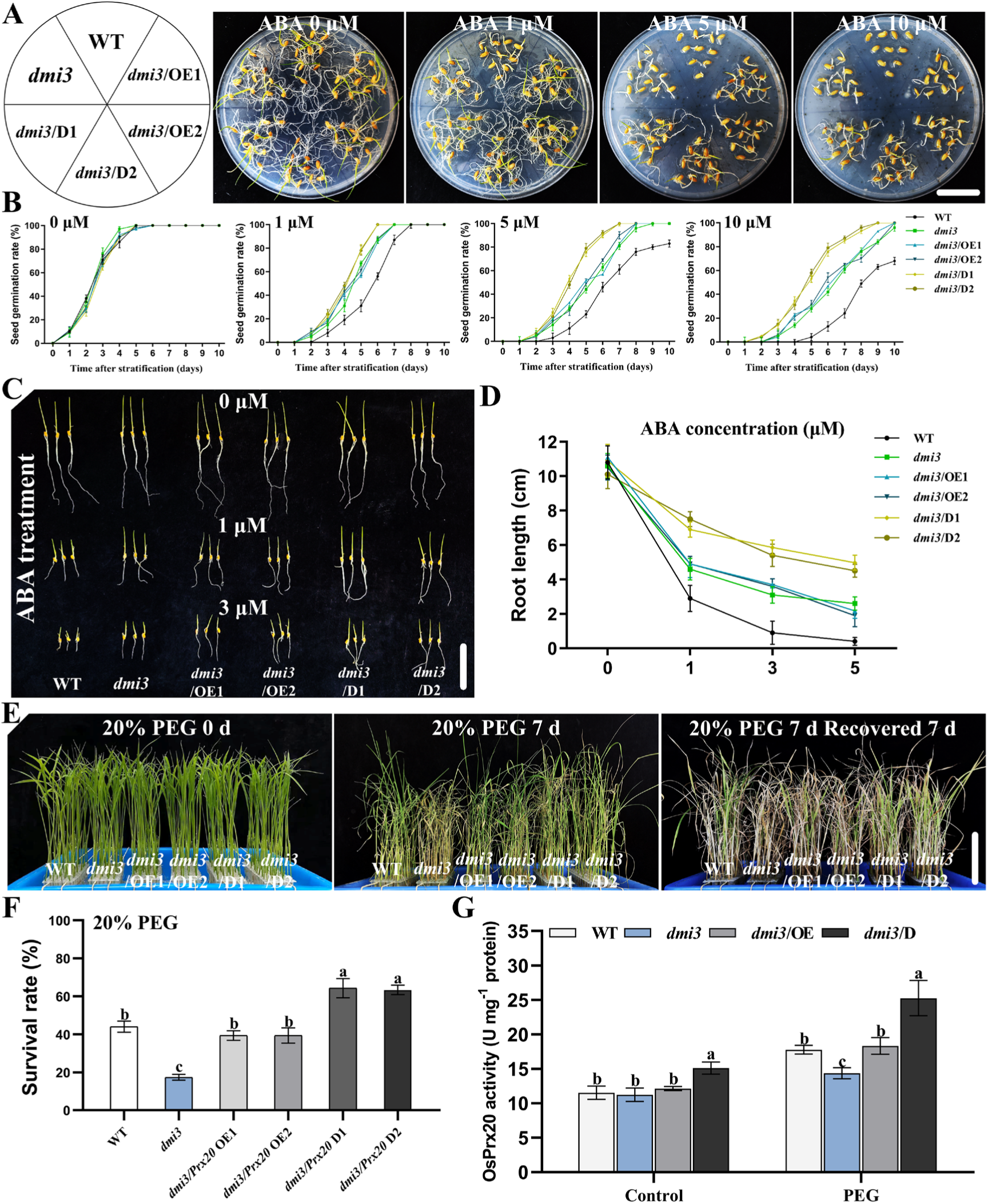
ABA-mediated OsDMI3-OsPrx20 pathway positively regulates osmotic stress tolerance. **(A)** Seed germination phenotypes of *osdmi3*-KO, *osdmi3/OsPrx20*-OE, *osdmi3/OsPrx20^T244D^*-OE, and WT were photographed after treatment with different concentrations of ABA (0, 1, 5, and 10 μM) for 10 d. Scale bar, 3 cm. **(B)** The seed germination rates are shown in (A). **(C)** Primary root growth phenotypes of *osdmi3*-KO, *osdmi3/OsPrx20*-OE, *osdmi3/OsPrx20^T244D^*-OE, and WT were photographed after treatment with different concentrations of ABA (0, 1, and 3 μM) for 5 d. Scale bar, 5 cm. **(D)** Primary root lengths of the materials shown in (C) grown in different concentrations of ABA as indicated for 10 d. **(E)** Phenotypes of *osdmi3*-KO, *osdmi3/OsPrx20*-OE, *osdmi3/OsPrx20^T244D^*-OE, and WT exposed to osmotic stress. 15-day-old rice seedlings were treated in 20% PEG-6000 solution for 7 d and then recovered by re-watering for 7 d. Scale bars, 10 cm. **(F)** The survival rate (%) of *osdmi3*-KO, *osdmi3/OsPrx20*-OE, *osdmi3/OsPrx20^T244D^*-OE, and WT after recovery by re-watering for 7 d is shown in (E). **(G)** The OsPrx20 activity in *osdmi3*-KO, *osdmi3/OsPrx20*-OE, *osdmi3/OsPrx20^T244D^*-OE, and WT after treated in 20% PEG-6000 solution for 2 d. Assay OsPrx20 activity using the Amplex® Red Hydrogen Peroxide/Peroxidase Assay Kit. The peroxidase standard curve was made with the HRP standard. The results of all experiments were repeated at least three times with similar results. Approximately 48 seedlings of each rice material were used per replicate in (E). Data in (B, F, and G) are means ± SD (n = 3). Data in (D) are means ± SD (n = 12). Significant differences are indicated: different lowercase letters, *P* < 0.05 (one-way ANOVA and Tukey’s multiple comparison tests).

To further confirm that the OsDMI3-OsPrx20 pathway can regulate the tolerance of rice plants to osmotic stress, *osdmi3*-KO, *osdmi3/OsPrx20*-OE, *osdmi3/OsPrx20^T244D^*-OE, and WT seedlings were treated with PEG, and the phenotypes and physiological indicators of these materials under stress conditions were analyzed. Compared with *osdmi3*-KO, *osdmi3/OsPrx20*-OE and *osdmi3/OsPrx20^T244D^*-OE exhibited a markedly increased tolerance to osmotic stress **(Fig. 6E and 6F)**, with a higher survival rate **(Fig. 6G and 6H)**, lower oxidative damage **(Fig. S12)**, and higher level of OsPrx20 activity **(Fig. 6E)**, in which *osdmi3/OsPrx20^T244D^*-OE showed better tolerance to osmotic stress than *osdmi3/OsPrx20*-OE. These results indicate that OsDMI3-mediated OsPrx20 Thr-244 phosphorylation plays an essential role in the tolerance of rice plants to osmotic stress. However, it should be noted that the tolerance of *osdmi3/OsPrx20*-OE to osmotic stress was similar to that of WT rather than that of *osdmi3*-KO, suggesting that under osmotic stress, in addition to the OsDMI3-OsPrx20 pathway, there is also OsDMI3-independent OsPrx20 activation pathway. Finally, we also compared the lignin content among *OsPrx20*-OE, *OsPrx20^T244A^*-OE, *OsPrx20^T244D^*-OE, *osdmi3/OsPrx20*-OE, and *osdmi3/OsPrx20^T244D^*-OE, and no variation was found in the different tissues of either seedlings or mature plants **(Fig. S13)**. These results indicate that the OsDMI3-OsPrx20 pathway is not involved in the regulation of lignin content in rice plants.

## 3. Discussion

Intracellular ROS as a signaling molecule requires maintaining at a low level. Even as an immunity molecule requires a relatively high level, it is also ephemeral. Its typical producers, RBOHs, generate widely varying levels of apoplastic H_2_O_2_ **(Kaya et al., 2019)**, which leads to the fact that lack of *OsRbohB/E* reduces stress tolerance, whereas lack of *OsRbohD/H* increases under the same stress in rice **(Shen et al., 2023)**. ROS bursts mainly occur at the site of attempted invasion in the early stages of plant-pathogen interactions. However, the second ROS bursts also occur during long-term infection, which triggers a hypersensitive reaction (HR), causing cell death. Interestingly, the second ROS bursts disappear in symbiotic interactions **(Nanda et al., 2010)**, considering important roles of ROS trade-off between early immunity and long-term defense in plants. Currently, the functions of Prxs under abiotic or biotic stress are still mainly focused on the scavenging of ROS, such as overexpression of *AtPrx22/AtPrx39/AtPrx69* in Arabidopsis, *TaPrx-2A* in wheat or *IbPrx17* in sweet potato positively regulates cold, salt, or drought tolerance by scavenging ROS **(Kim et al., 2012; Su et al., 2020; Zhang et al., 2022)**, which are consistent with overexpression of *OsPrx20* **(Fig. 2A, S4A and 5F)**. Also, inhibition of *LePrx06* in tomato increases H_2_O_2_ accumulation to enhance disease resistance **(Coego et al., 2005)**, as knockout of *OsPrx20* **(Fig. 2C and S8A)**. Under biotic stress, however, it was also shown that down-regulation of *AtPrx33/AtPrx34* in Arabidopsis or silencing of *CaPrx2* in pepper decreases ROS levels to increase susceptibility to pathogens **(Choi et al., 2007; Daudi et al., 2012),** implying Class III Prxs have different effects in abiotic or biotic stress. Both OsPrx30 and OsPrx114, belonging to the rice Class III Prxs, scavenge ROS generated under stress by increasing POD activity; overexpression of *OsPrx114* enhances drought tolerance, whereas overexpression of *OsPrx30* reduces bacterial blight resistance **(Liu et al., 2021b; Zheng et al., 2023)**, directly implying that Class III Prxs plays different roles in maintaining ROS homeostasis under different stress conditions. These functions were further demonstrated in OsPrx20. Overexpression of *OsPrx20* enhanced osmotic stress tolerance through increasing POD activity to scavenge excess ROS generated **(Fig. 2A and S4B)**, but it was also more sensitive to rice blast **(Fig. 2C)**. The bidirectional regulation of OsPrx20 under different stresses fully indicates its essential role in regulating ROS homeostasis. Meanwhile, ABA-mediated OsPrx20 Thr-244 phosphorylation by OsDMI3 **(Fig. 5F)** further enhances osmotic stress tolerance **(Fig. 4C and 4D)**.

ABA induces the production of extracellular H_2_O_2_ and its entry into the cytosol to form intracellular ROS signals, and extracellular ROS production is also accompanied by intracellular ROS clearing, thus maintaining the dynamic balance of intracellular ROS levels. However, ABA-induced and PEG-induced phosphorylation of the Thr-244 site of OsPrx20 did not appear in o*sdmi3*-KO **(Fig. 5E and 5G)**, suggesting OsPrx20 Thr-244 phosphorylation was directly dependent on OsDMI3. Meanwhile, by comparing the intracellular ROS signals of different material protoplasts under ABA treatment, the highest levels of intracellular ROS in *osprx20*-KO and the lowest in *OsPrx20^T244D^*-OE were found **(Fig. 5H)**, suggesting OsPrx20 has an ability to scavenge ABA-induced intracellular ROS and its Thr-244 phosphorylation further enhances this ability. Interestingly, the OsPrx20 Thr-244 phosphorylation, while enhancing its ROS scavenging capacity, further reduces the resistance to rice blast **(Fig. 4H)**. ABA also regulates seed germination and root growth. Lack of *OsDMI3*, as Ca^2+^ signal decoder, decreased sensitivity to ABA **(Ni et al., 2019)**. Lack of *OsRbohB*, as ROS signal producer, also reduced the sensitivity to ABA **(Wang et al., 2023)**. However, lack of *OsPrx20* enhanced sensitivity to ABA **(Fig. S11)**. OsPrx20 Thr-244 phosphorylation further relieved the inhibitory effect of ABA on seed germination and root growth **(Fig. 5A and 5C)**, indicating that phosphorylation of the OsPrx20 Thr-244 is crucial in responding to ABA, but its role is not to amplify or transmit signal. In the background of *osdmi3*-KO, the sensitivity to ABA of *osdmi3*/*OsPrx20*-OE was basically the same as that of *osdmi3*-KO, whereas *osdmi3*/*OsPrx20^T244D^*-OE by ABA-inhibited was significantly better than that of *osdmi3*-KO **(Fig. 6A and 6C)**, suggesting OsPrx20 acts downstream of OsDMI3 and involves in ABA-regulated seed germination and root growth by OsDMI3 phosphorylating it at Thr-244. OsDMI3 integrates and amplifies Ca^2+^ and H_2_O_2_ signaling in ABA-mediated abiotic stress response to external environmental stress through phosphorylation downstream **(Torres and Berlanga, 2023)**, such as ABA-mediated phosphorylation of Ser-191 in OsRbohB and Thr-25 in OsMKK1 by OsDMI3 **(Chen et al., 2021; Wang et al., 2023)**. Coincidentally, unlike OsDMI3 phosphorylation of OsRbohB that promotes H_2_O_2_ production to amplify ROS signaling, phosphorylation of OsPrx20 directly enhanced its activity to promote the scavenging of intracellular ROS **(Fig. 5H)**, which was somewhat analogous to the phosphorylation of OsUXS3 by activated-OsDMI3 to enhance OsCATB activity and thereby scavenging ROS **(Ni et al., 2022)**, but more directly. Remarkably, ABA produced under stress conditions enhanced response to abiotic stress, but it also inhibits growth and development. However, this limitation has broken in OsPrx20. Overexpression of *OsPxr20* enhanced osmotic stress tolerance while enlarging spike and grain size **(Fig. 3D and 3G)** and increasing grain number per spike **(Fig. 3E)** without affecting plant height, tiller number, and seed setting rate **(Fig. 3B, 3C, and 3F)**. Nevertheless, combined with the susceptibility of *OsPrx20*-OE to rice blast, the elevated yield of a single rice after overexpression of *OsPrx20* alone could not be regarded as a perfect excellent breeding trait.

The growth phenotype of a single knockout of *OsPxr20* reduced plant height **(Fig. 3A)** but unaffected tiller number and seed setting rate **(Fig. 3C and 3F)**, consistent with the Green Revolution as marked by the breeding of dwarfism varieties **(Hedden, 2003)**. However, lack of *OsPxr20* also caused by smaller single spike size **(Fig. 3D)** and fewer grain number per spike **(Fig. 3E)**. Although it unaffected seeds smaller **(Fig. 3I, 3J, and 3K)**, it affected the yield of a single rice. OsPrx20 was mainly localized in the cytoplasm rather than the cell wall **(Fig. S1C and S1D)**. Class III Prxs have been reported to promote lignin synthesis **(Cosio and Dunand, 2009; Shigeto and Tsutsumi, 2016)**, but it doesn’t mean they can only play a role in the cell wall. Indeed, monolignols and enzymes are unlikely to interact directly due to the limited freedom of movement in the cell wall during growth **(Vanholme et al., 2010)**. The *ρ*-coumaric acid (*ρ*CA) is an excellent substrate for peroxidase in grasses. It has been proposed as a radical shuttle for lignification, efficiently transferring its unpaired electron to sinapyl alcohol and lignin polymers **(Hatfield et al., 2008)**. Lignin, one of the main components of the cell wall, is vital to plant cells’ morphology, tissue structure, and against environmental stresses and pests. Lignin-deficient plants are enormously impaired in their development, most showing dwarfism **(Bonawitz et al., 2014)**, as shown in *osprx20*-KO **(Fig. 3A and S4B)**. However, overexpression of *OsPrx20* significantly increased lignin content in the roots and stems of mature plants **(Fig. S4)** but did not affect plant height **(Fig. 3B)**. Then, it is debatable whether the dwarfism of *osprx20*-KO is directly due to impaired lignin synthesis because lignification, particularly if ectopic, may also constrain growth **(Ha et al., 2021)**. Meanwhile, OsPrx20, expressed with the highest expression in stems and the lowest in leaves but not in root tips and root meristem **(Fig. S1A and S1B)**, might act indirectly on lignin synthesis in the more lignified xylem cells in stems and roots while increasing xylem strength. Lignin deposition is crucial for stem bending resistance, directly determining the rice’s ability to resist lodging. Still, this aspect has not been further explored in the present study. A previous study reported that overexpression of *CsPrx25* enhances lignification as an apoplastic barrier for *Xanthomonas citri subsp. citri* (*Xcc*) infection in citrus **(Li et al., 2020)**. However, it is clear that while overexpression of *OsPrx20* increased lignin content, it provided insufficient disease resistance. A previous study has directly demonstrated that the transcriptional level of two rice Class III Prx genes (*Os05g04470* and *Os10g39170*) under rice blast infection was repressed **(Li et al., 2017)**. Similarly, the expression of *OsPrx20* and the levels of OsPrx20 also showed significant down-regulation under rice blast infection **(Fig. 2F and 2G)**, probably through transcriptional regulation to represses OsPrx20 to maintain a relatively high level of ROS to protect against rice blast, which provides a direction that can be considered for subsequent in-depth research. In addition, A previous study also has found that semi-dwarf crops typically have more excellent resistance to various diseases and insects **(Liu et al., 2020)**, as in *osprx20*-KO **(Fig. 2C)**. Combining all phenotypes, however, it is not always feasible to pursue the development of dwarfism and high-yielding rice varieties by changing a single gene (e.g., knockout or overexpression of *OsPrx20*). It is even more essential to balance growth and stress tolerance — better adaptation to external environments while achieving stable yield increases in rice.

The key of OsDMI3 as a Ca^2+^/CaM-dependent protein kinase in response to adversity stress comes from stress-generated ABA and Ca^2+^ signals, and its current involvement in a range of cellular processes is dependent on ABA **(Chen et al., 2021; Ni et al., 2019; Shi et al., 2014; Wang et al., 2023)**. However, during normal growth, ABA and intracellular Ca^2+^ concentrations derived from stimuli generated by external environmental changes are low, which may result in OsDMI3 inactivating. The current reports involved in regulating growth and development are also mainly mediated by jasmonic acid (JA), gibberellins (GA), brassinosteroids (BRs), and auxin **(Duan et al., 2023; Jin et al., 2023; Qi et al., 2023)**, whereas it is unknown whether OsDMI3 responds to these signals. No significance of the grain traits in *OsPxr20*-OE, *OsPrx20^T244A^*-OE, and *OsPrx20^T244D^-*OE **(Fig. S9)** and the lignin content in *OsPxr20*-OE, *OsPrx20^T244A^*-OE, *OsPrx20^T244D^*-OE *osdmi3*/*OsPxr20*-OE, and *osdmi3*/*OsPrx20^T244D^*-OE under normal conditions **(Fig. S10 and S13)**, suggesting that OsDMI3-OsPrx20 pathway did not regulate grain development and lignin synthesis. Although OsPrx20 activity was increased by phosphorylation at the Thr-244 site under normal conditions **(Fig. 4E and 6G)**, grain fullness and lignin synthesis did not further increase, which may be limited by the concentration of biosynthetic substrates. However, the OsDMI3-mediated phosphorylation of Thr-244 or other sites in OsPrx20 under adverse growth conditions, whether directly involved in regulating lignin synthesis and development, is unknown.

In brief, regulation of ROS homeostasis through OsPrx20 is key to the trade-off between defense in abiotic and immunity in biotic in rice, and it also participates in growth and development. On the one hand, OsPrx20 responds to ABA signaling and positively regulates osmotic stress defense by phosphorylating the Thr-244 in OsPrx20 by OsDMI3. Lack of *OsPrx20* directly weakens cells’ ability to scavenge ROS under osmotic stress. On the other hand, OsPrx20 is a negative regulator of resistance to rice blast. Lack of *OsPrx20* enhances the immunity to rice blast, whereas overexpression of *OsPrx20* is more sensitive. Meanwhile, lack of *OsPrx20* achieves dwarfism phenotype and smaller spikes under normal conditions, while overexpression of *OsPrx20* enlarges the spike size and grain fullness **(Fig. 7)**. Still, it is uncertain whether the changes in plant height, spike, and grain are caused by an imbalance of ROS homeostasis under normal conditions due to the deletion of *OsPrx20*, which leads to insufficient oxidation of substrates such as lignin synthesis precursors, or whether OsPrx20 can be directly involved in the regulation of genes related to growth and development. Also, this study does not delve into the potential regulatory mechanisms of the OsDMI3-OsPrx20 molecular module on growth and development under stress, providing an exciting future research direction.

**Figure 7.**
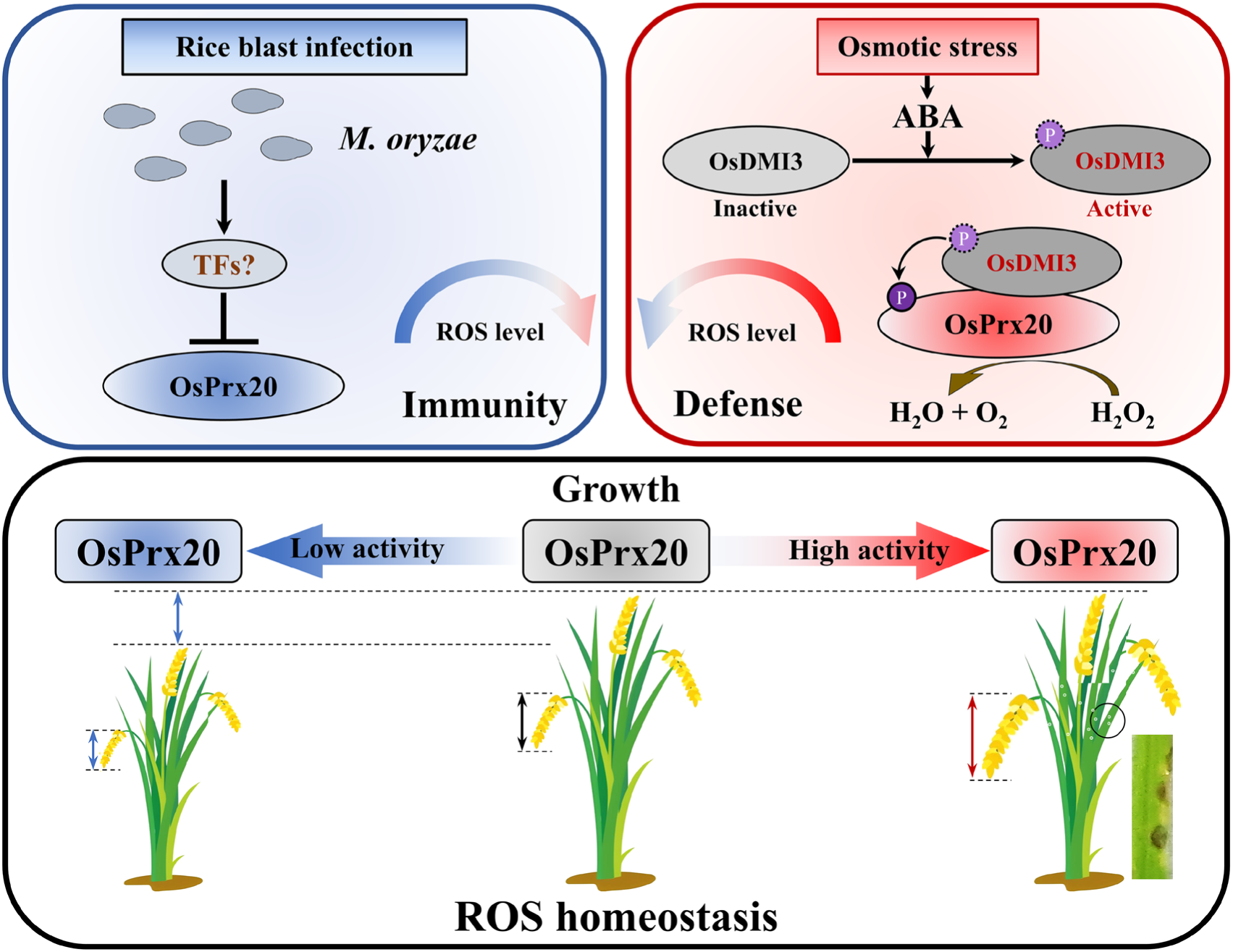
A working model of OsPrx20 in regulating stress response and growth in rice. **(Left)** Under rice blast infection, the depression and down-regulation of *OsPrx20* increase the ROS levels to enhance the immunity to rice blast. **(Right)** Under osmotic stress, ABA-mediated OsDMI3 phosphorylate OsPrx20 Thr-244 enhances osmotic stress tolerance by scavenging intracellularly accumulated ROS. **(Below)** Under normal conditions, OsPrx20 maintains intracellular ROS homeostasis. Lack of *OsPrx20* leads to dwarfism and a smaller spike; overexpression of *OsPrx20* enlarges the spike and grain size but weakens resistance to disease infection. Blunt-ended arrows indicate repression, and solid arrows indicate the activity process.

## 4. Materials and Methods

### 4.1 Rice materials

Rice (*Oryza sativa* L.) used in this study include *osprx20*-KO, *OsPrx20*-OE, *OsPrx20^T244A^*-OE, *OsPrx20^T244D^*-OE, *osdmi3*-KO, *osdmi3/OsPrx20*-OE, *osdmi3/OsPrx20^T244D^*-OE, *proOsPrx20*-GUS, and wild type (*O. sativa* sub. *japonica* cv Nipponbare; WT) Among them, *OsPrx20*-OE, *OsPrx20^T244A^*-OE, *OsPrx20^T244D^*-OE, or *proOsPrx20*-GUS transgenic plants via Agrobacterium (*Agrobacterium tumefaciens*)-mediated transformation of the pCambia1304-*OsPrx20*, pCambia1304-*OsPrx20^T244A^*, pCambia1304-*OsPrx20^T244D^*, or pCXGUS-*proOsPrx20(-2111 bp* - *-1 bp)* construct were transformed into WT, respectively. *osdmi3*-KO were described in previous studies **(Ni et al., 2019)**. *osdmi3*/*OsPrx20*-OE or *osdmi3*/*OsPrx20^T244D^*-OE transgenic plants via Agrobacterium-mediated transformation of the pCambia1304-*OsPrx20* or pCambia1304-*OsPrx20^T244D^* construct were transformed into *osdmi3*-KO, respectively. *osprx20*-KO by CRISPR/Cas9 edited gene mutants and identification results were shown in Supplemental Figure **(Fig. S3A)**. All materials were selected on 50 μg/mL Hygromycin-B. The gene expression of each transgenic strain was also assayed using RT-qPCR, and the results were shown in Supplemental Figure **(Fig. S3B)**. The homozygous T2 or T3 seeds of mutants and transgenic plants were used for further analysis. The rice seedlings’ growth and stress conditions are provided in **Supplemental Method S1**. The primers used for identifying mutants, RT-qPCR, and constructing recombination vectors were listed in **Supplemental Table S1-3.**

### 4.2 Pathogen Inoculation

*M. oryzae* (Guy11) strain was used for rice infection. Guy11 strain was first inoculated on a complete medium (CM) that grew at 26℃ under a 12 h/12 h (light/dark) condition. Spores were collected 2 weeks after inoculation and resuspended to 1 × 10^5^ spores/mL, then spray-inoculated on three-leaf-stage rice seedlings. Inoculated plants were kept in a growth chamber at 26°C and 90% humidity in the dark for the first 24 h, followed by a 12 h/12 h (light/dark) cycle for 5 days. Disease symptoms were examined 5 days later. For fungal biomass assays, genomic DNA extracted from inoculated leaves was used for Real-time quantitative PCR (RT-qPCR) in **Supplemental Method S2**. Relative fungal biomass was calculated as the amount of *Pot2* in *M. oryzae* relative to *OsUbiquitin* in rice. The primers used for RT-qPCR were listed in **Supplemental Table S2.**

### 4.3 Phenotypic analysis

Treatments with ABA to determine the seed germination and primary root growth were performed as described previously **(Chen et al., 2021)**. Treatments with 20% PEG-6000 solution to determine the stress phenotype were performed. Firstly, seedlings with the same growth were pretreated in nutrient solution for 2 days to eliminate damage to the roots. For PEG-6000 treatments, the PEG was gradually added to the nutrient solution during PEG treatment and finally reached a 20% (w/v) concentration. For ABA treatments, rice seedlings need shielding to minimize exposure to light.

### 4.4 Immunoblotting analysis

Protein extraction buffer and immunoblot analysis as described previously **(Ni et al., 2019)**. The target proteins were separated by 12% or 15% (w/v) SDS-PAGE. Antibodies used in this study include anti-GST antibody (AE001; ABconal; Wuhan, China; 1:5,000, v/v), anti-His antibody (AE003; ABconal; Wuhan, China; 1:5,000, v/v), anti-Actin (plant specific) antibody (AC009; ABconal; Wuhan, China; 1:5,000, v/v), anti-pThr antibody (AP1422; ABconal; Wuhan, China; 1:1,000, v/v), anti-phospho-OsPrx20^Thr244^ antibody (ABconal; Wuhan, China; 1:1,000, v/v), anti-OsDMI3 antibody **(Wang et al., 2023)**, anti-OsPrx20 antibody (ABconal; Wuhan, China; 1:1,000, v/v), and Goat anti-Mouse/Rabbit IgG-HRP antibody (M21003; Abmart; Shanghai, China, 1:10,000, v/v). The specificity of the anti-OsPrx20 antibody is shown in **Supplemental Figure (Fig. S3C)**.

### 4.5 Agronomic traits analysis

To evaluate development components of different materials under the field. During the tillering stage, different materials were selected to determine their plant height and tiller number. During the maturity stage, phenotype, grain number, and seed setting rate of single-spike in different materials were counted. Grain length, grain width, and grain thickness were measured using a vernier caliper, and the thousand kernel weight was quantified. The data was statistically analyzed via one-way ANOVA and Tukey’s multiple comparison tests using SPSS (Version 21.0) software.

### 4.6 Protein-protein interaction assay

In Y2H assay, the coding sequence (CDS) of *OsPrx20* or *OsDMI3* was inserted into pGADT7 (AD) or pGBKT7 (BD) vector, respectively. The AD and BD recombinant plasmids were transformed into yeast Y2HGold strain together. In GST pull-down assay, the CDS of *OsPrx20* or *OsDMI3* was inserted into pET-30a or pGEX4T-1 vector, respectively. OsPrx20-His and OsDMI3-GST were expressed in *Escherichia coli* (BL21). OsDMI3-GST or GST alone was immobilized on GST magnetic beads and incubated with OsPrx20-His in a binding buffer at 4°C for 2 h. The particles were washed with wash buffer and boiled in 1× SDS loading buffer. Samples were separated on 12% SDS-PAGE gels and analyzed by IB using an anti-GST antibody or anti-His antibody, as described previously **(Ni et al., 2019)**. In LCI assay, the CDS of *OsPrx20* or *OsDMI3* was inserted into pCAMBIA1300-nLUC or pCAMBIA1300-cLUC vector, respectively. The constructs were co-transfer infiltrated in tobacco leaves by the *Agrobacterium*-mediated method. OsPP45-nLUC/OsDMI3-cLUC were used as a positive control **(Ni et al., 2019)**. In BiFC assay, the CDS of *OsPrx20* or *OsDMI3* was inserted into pSPYNE or pSPYCE vector. Constructs expressing OsPrx20-YFP_N_ and OsDMI3-YFP_C_ were co-transfected into rice protoplasts using the PEG-mediated method. More details of experiments are provided as described previously and in **Supplemental Methods S3-5**. The primers used for constructing recombination vectors are listed in **Supplemental Table S3**.

### 4.7 LC-MS/MS analysis

OsPrx20-His was incubated with OsDMI3-GST in kinase reaction solution (25 mM Tris-HCl [pH 7.5], 5 mM MnCl_2_, 0.5 mM CaCl_2_, 2 μM CaM, 1 mM DTT, 10 μM ATP) at 30°C for 30 min. Then, using trypsin (10 ng/μl, V5113, Promega) to digest the phosphorylated OsPrx20-His and analyzed by the LC-MS/MS system (LTQ Orbitrap XL ETD; ThermoFisher Scientific, Waltham, MA, USA) as described previously **(Gampala et al., 2007)**. Proteome Discoverer software (1.2 version, ThermoFisher Scientific; Waltham, MA, USA) was used to analyze the phosphorylated site.

### 4.8 In vitro phosphorylation assay

5 μg kinase OsDMI3-GST was incubated with 5 μg substrates (OsPrx20-His, Os Prx20^T244A^-His, or myelin basic protein [MBP; Sigma-Aldrich]) in kinase reaction solution for 30 min as described previously **(Shen et al., 2023b)**. 5×SDS-PAGE loading buffer was added to terminate the reaction, and the reaction mixtures were separated by 15% SDS-PAGE. Phosphorylated substrates were detected by autoradiography using a storage fluorescent screen (Typhoon TRIO; Amersham Biosciences, Piscataway, NJ, USA).

### 4.9 In vivo phosphorylation assay

OsPrx20 proteins were immunoprecipitated from the total protein extracted from WT or *osdmi3*-KO after ABA-treated by an anti-OsPrx20 antibody bound to protein A/G beads and separated by 12% SDS-PAGE. The phosphorylated OsPrx20 was detected by an anti-pThr antibody, and the total OsPrx20 protein was detected using an anti-OsPrx20 antibody. The chemiluminescence was captured using the Tanon-5200 image system (Tanon; Shanghai, China).

### 4.10 OsPrx20 activity assay

OsPrx20 proteins were first immunoprecipitated from the total protein extracted from different materials after PEG treatment by an anti-OsPrx20 antibody bound to protein A/G beads. Then, an Amplex^®^ Red Hydrogen Peroxide/Peroxidase Assay Kit (A22188; ThermoFisher Scientific; Waltham, MA, USA) was used to assay the OsPrx20 activity according to the manufacturer’s instructions. Fluorescence was measured with a SpectraMax iD5 Microplate Reader (Molecular Devices; Sunnyvale, CA, USA) using excitation at 530 nm and fluorescence detection at 590 nm. The OsPrx20 activity in each sample was calculated using a standard curve. The peroxidase standard curve was made with the horseradish peroxidase (HRP) standard.

### 4.11 Fluorescence probe for ROS detection

CM-H_2_DCFDA (C6827; ThermoFisher Scientific, Waltham, MA, USA) was used as a probe to detect ROS in rice protoplasts after rapid treatment with 1 μM ABA. Using a scanning confocal microscope (LSM880; Carl Zeiss; Oberkochen, Germany), excitation at 450-490 nm and emission at 515-565 nm were observed as described previously **(Kuběnová et al., 2023)**.

## Accession Numbers

Sequence data in this article can be found in the following accession numbers: *OsPrx20*, LOC_Os01g73170; *OsDMI3*, LOC_Os05g41090; *OsGAPDH,* LOC_Os02g38920; *OsUbiquitin,* LOC_Os03g13170 (Rice Genome Annotation Project database); *M. oryzae Pot2*, Z33638 (National Center for Biotechnology Information).

## Acknowledgments

This work was supported by the National Natural Science Foundation of China (Grant No. 31971824, 32170316, and 32372038).

## Author Contributions

L.N. and M.J. conceived the project and designed the experiments. T.S. performed most of the experiments and analyzed the data. W.Q., D.C., J.H., R.Y., X.J., F.D., B.X., Z.H., H.J., and Z.Z. performed some experiments. T.S. wrote the manuscript. L.N. and M.J. revised the manuscript.

## Data availability

All data supporting this study’s findings are available in the main text or the Supporting Information. All sequence data can be found in the Rice Genome Annotation Project (http://rice.plantbiology.msu.edu/) and the National Center for Biotechnology Information (https://www.ncbi.nlm.nih.gov/).

## Supporting Information

Additional Supporting Information may be found online in the Supporting Information section at the end of the article.

### 1. Supporting Figures

**Figure S1.** Expression analysis and subcellular localization of OsPrx20.

**Figure S2.** Protein levels of each transformed gene in LCI and BiFC assays.

**Figure S3.** Identification of knockout mutants, overexpression transgenic rice, and specificity of the anti-OsPrx20 antibody.

**Figure S4.** Indicators of oxidative damage in *osprx20*-KO, *OsPrx20*-OE, and WT under osmotic stress.

**Figure S5.** Lignin content of *osprx20*-KO, *OsPrx20*-OE, and WT.

**Figure S6**. Indicators of oxidative damage in *osprx20*-KO, *OsPrx20*-OE, *OsPrx20^T244A^*-OE, *OsPrx20^T244D^*-OE, and WT under osmotic stress.

**Figure S7.** OsPrx20 activity of OsPrx20-His, OsPrx20^T244A^-His, OsPrx20^T244D^-His, or His expressed in vitro.

**Figure S8.** H_2_O_2_ content and DAB staining in the leaves of *osprx20*-KO, *OsPrx20*-OE, *OsPrx20^T244A^*-OE, *OsPrx20^T244D^*-OE, and WT after infected with Guy11.

**Figure S9.** Phenotypic analyses of *osprx20*-KO, *OsPrx20*-OE, *OsPrx20^T244A^*-OE, *OsPrx20^T244D^*-OE, and WT spikes and grains.

**Figure S10.** Lignin content of *osprx20*-KO, *OsPrx20*-OE, *OsPrx20^T244A^*-OE, *OsPrx20^T244D^*-OE, and WT.

**Figure S11.** Seed germination phenotypes of *osprx20-KO*, *OsPrx20*-OE, and WT.

**Figure S12.** Indicators of oxidative damage in *osdmi3*-KO, *osdmi3/OsPrx20*-OE, *osdmi3/OsPrx20^T244D^*-OE, and WT under osmotic stress.

**Figure S13.** Lignin content of *OsPrx20*-OE, *OsPrx20^T244A^*-OE, *OsPrx20^T244D^*-OE, *osdmi3/OsPrx20*-OE and *osdmi3/OsPrx20^T244D^*-OE.

### 2. Supporting Methods

**Method S1.** Rice seedling growth and stress conditions

**Method S2.** RT-qPCR

**Method S3.** Y2H assay

**Method S4.** LCI assay

**Method S5.** BiFC assay

**Method S6.** GUS (β-glucuronidase) staining

**Method S7.** Subcellular localization

**Method S8.** H_2_O_2_ content

**Method S9.** Total POD activity

**Method S10.** DAB staining

**Method S11.** MDA content

**Method S12.** Electrolyte leakage rate

**Method S13.** Lignin content

### 3. Supporting Tables

**Table S1.** PCR primers are used for identifying mutants.

**Table S2.** PCR primers are used for RT-qPCR.

**Table S3.** PCR primers are used for constructing recombination vectors.

**Table S4.** PCR primers are used for cloning genes and promoters.

